# A Statistical Guideline for Analyzing Repeated-Measures Proteomics Data

**DOI:** 10.64898/2026.09.18.752572

**Authors:** Mikkel Skjoldan Svenningsen, Julie Lyng Forman, Alicia Lundby

## Abstract

Mass spectrometry-based proteomics increasingly supports study designs with repeated measurements, such as longitudinal sampling and spatial tissue profiling. These designs offer rich biological insights but introduce statistical challenges due to within-subject correlations and variance heterogeneity across time points or anatomical locations. Despite the growing use, best practices for analyzing repeated-measures proteomics data remain limited. We systematically compare commonly applied methods, including paired t-tests and linear mixed-effects models with correlation induced by random effects, to more flexible covariance-pattern models with unstructured covariance. Through simulation studies, we evaluate false-positive rates and statistical power under varying conditions of covariance heterogeneity. Our results demonstrate substantial limitations of conventional approaches and highlight the robustness and superior performance of unstructured covariance pattern models for repeated measures proteomics. This approach accommodates diverse study designs supporting accurate inference and reproducibility. We demonstrate that conventional F-tests can be severely biased, not just for small sample sizes but also moderately large ones, while using pairwise t-tests or applying parametric bootstrapping to evaluate the p-value of the F-tests mitigates the problem. We provide real world data examples showing how to perform valid and informative statistical analyses for temporal and spatial proteomics studies with repeated measurements.

## 1. Introduction

Mass spectrometry-based proteomics has transformed the ability to characterize deep proteomes in both tissues and biofluids. Continuous improvements in instrument sensitivity, resolution and quantitative accuracy now enable robust proteome profiling from minimal sample input[1-3]. These advances, coupled with innovations in acquisition speed and automation, are driving the field towards high-throughput workflows [4-6], enabling study designs with larger cohorts and increased biological replication. High-throughput capability further facilitates repeated measurements within individuals, enabling longitudinal tracking of proteome dynamics [7-9] and spatial heterogeneity assessment within tissues [10-14]. Repeated measures designs involve sampling multiple protein intensities from the same experimental unit, such as a participant, an animal or cell culture, across different time points or anatomical locations. These designs violate the independence assumption underlying classical statistical methods, such as ordinary linear regression and ANOVA, and therefore necessitate specialized approaches that model correlation among repeated measurements.

The main objective of repeated measures proteomics analysis is to identify proteins whose abundance changes significantly over time, varies across tissue regions, or exhibit distinct temporal or spatial patterns between groups. This typically involves analyzing the protein intensity as the outcome variable, with time/location and group as covariates, and performing analyses separately for each protein. An effective approach should maximize statistical power to detect true differences while controlling false discovery rate, ensuring reliable identification of biologically meaningful patterns.

Current approaches often rely on linear mixed models with correlation induced by random effects. However, these models impose restrictive assumptions on within-subject correlations and neglect variance heterogeneity across time points or anatomical locations [15]. Paired t-tests are also commonly used but do not allow adjustment for additional covariates, limiting their applicability in complex study designs.

We propose covariance pattern models as a more robust framework for analyzing repeated measures proteomics. Both random effects models and covariance pattern models are linear mixed models, but they differ in how they specify the covariance structure. Covariance pattern models model correlations and variances directly, offering greater flexibility than random effects models. Misspecifying the covariance structure can substantially affect inference, leading to incorrect identification of time-varying or location-dependent proteins. Another challenge is statistical testing in small to moderate size samples, where conventional F-tests can produce biased p-values. To address this, we propose using either pairwise t-tests or a parametric bootstrap version of the F-tests to ensure valid inference with limited sample size.

This study provides statistical guidance for analyzing repeated proteome measurements by building on and extending previous work in this field [16-19]. Our focus is on balanced study designs, including but not limited to proteomes measured from the same participants across multiple time points or distinct tissue regions. Our evaluation encompasses simulation-based assessments of false-positive rates and statistical power, as well as applications to real-world proteomics datasets using time and anatomical location as illustrative examples. Our findings support linear mixed models with an unstructured covariance pattern as the preferred approach for repeated measures proteomics data, owing to their flexibility across study designs. By clarifying the implications of statistical modeling choices, this study aims to promote robust and reproducible analysis of temporal and spatial proteomics data across diverse biological contexts. All R code used for simulations and analyses is provided as a supplementary material.

## 2. Methods

Repeated measures proteomics require statistical models that account for correlation among observations from the same subject. Selecting an appropriate analysis strategy involves several considerations. Critical aspects include:

i. Fixed effects: modeling population level effects of covariates
ii. Covariance structure: modeling variance and correlation between measurements
iii. Hypothesis testing: T-tests or F-tests and choice of degrees of freedom method

We restrict our study to balanced study designs, where all subjects are measured at the same repeated points (e.g. time points or anatomical sites). Analysis of unbalanced designs, which are common in analysis of biobank samples, require case by case modeling decisions and is complicated by the inherent risk of bias due to informative sampling [20].

### Fixed Effects: Modeling Population Level Effects of Covariates

The primary goal of repeated measures proteomics analysis is usually to identify proteins that change with time, vary within tissue, or show different temporal or spatial patterns between groups of interest. Single-group evaluation of temporal or spatial patterns in protein intensity can be achieved by including time or site as a covariate (aka fixed effect) in a linear mixed model [15]. Time (or anatomical site) should be treated as a categorical variable to capture any trajectory shape. This is done to avoid imposing restrictive assumptions such as linearity, which can bias results if the true profile is non-linear. Figure 1 includes two examples of protein intensity profiles across four repeated measurements, to underscore the need for models that accommodate complex biological patterns. Group comparisons (e.g. diseased vs healthy) can be included in the analyses by adding group and the group^*^time interaction (or group^*^site for spatial analyses) in the model. Additional covariates, such as age or sex, can also be included for adjustment.

**Figure 1:**
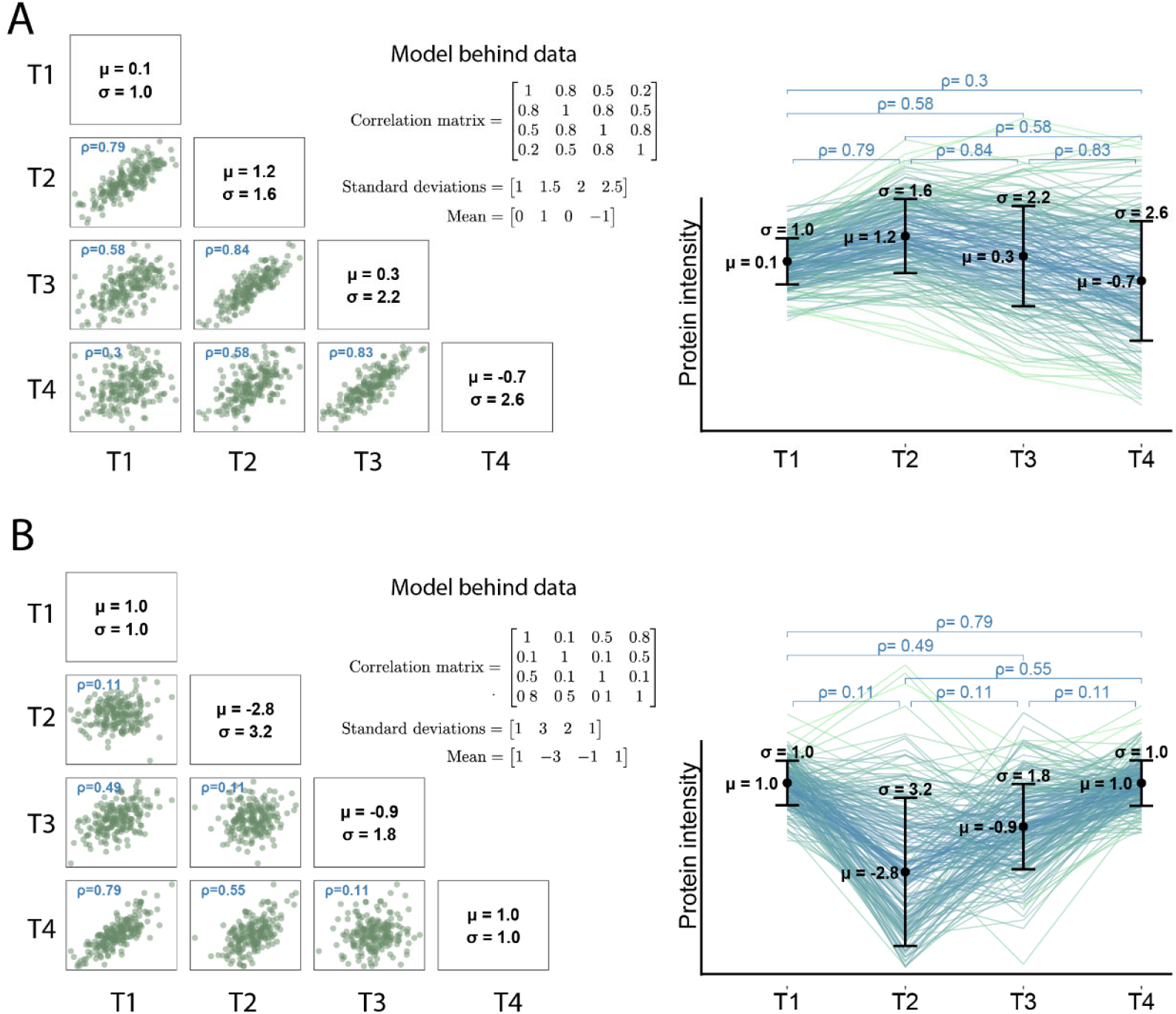
Response profiles and covariance patterns illustrated by two simulated repeated measures proteomics datasets with n=200 participants each. Mean values (µ), standard deviations (σ), and correlations (ρ) are shown for each scenario. Left panels show pairwise scatterplots of residuals between timepoints with estimated correlations. Right panels depict subject-specific trajectories with estimated means, standard deviations, and correlations between time points. A high positive correlation indicates that the same subject tends to be consistently above or below the population mean at both timepoints. The inset shows the mean and covariance parameters used to generate the data. (A) Dataset mimicking a typical clinical follow-up study with variance increasing over time and correlation decreasing with increasing time-distance. (B)Dataset mimicking a stimulus-experiment, where responses return to baseline as the stimulus effect wears off. Here, the strongest correlation occurs between baseline and the final measurement, which also share a lower variance compared to the intermediate timepoints.

### Covariance Structure: Modeling Correlation Between Measurements

Linear mixed models extend ordinary linear models by accounting for correlation among repeated measurements within the same subject as well as potentially time-or site-dependent variation. This increases the analytical complexity but has the advantage that differences in protein intensity can be detected with higher power, especially when the correlations are high [15]. In linear mixed models, correlations and variance between pairs of repeated measurements are summarized in a covariance matrix, which defines the assumed pattern of within-subject dependence (Figure 1).

To perform a repeated measures analysis, a particular model for the covariance structure must be chosen and different covariance structures impose different assumptions. If the chosen structure deviates from the true pattern, inference may suffer from inflated false positive rates and/or reduced power [15]. The simplest, and most commonly applied, models assume a covariance pattern called *compound symmetry*, which has a single correlation parameter and a single variance parameter shared across all measurement points (e.g. time or space). Compound symmetry is often modeled indirectly by including a random effect for each subject (a random intercept) in the linear mixed-model. As Figure 1 illustrates, the compound symmetry structure is often too restrictive compared to the patterns found in biological experiments, likely leading to faulty inference in many applications. The most flexible covariance pattern is the *unstructured covariance*, which has distinct correlations for each pair of repeated measurement points and a distinct variance for each measurement point. That is, unstructured covariance does not make any restrictions on the pattern. This flexibility makes the unstructured covariance the preferred choice for analysis of repeated measures proteomics data.

The downside to the unstructured covariance is the large number of parameters which need to be estimated. With *k* measurement points (e.g. time points or spatial sites), there are *k*(*k* − 1)/2correlations + *k* variances. This can lead to convergence issues when the number of study participants is small [21]. A commonly raised concern about the unstructured covariance model is that the high number of parameters lowers the statistical power compared to simpler covariance patterns such as compound symmetry. However, our in silico study only showed a diminutive power loss in case the true covariance pattern was a compound symmetry pattern. We only found substantial differences in power in cases where the true pattern deviated from compound symmetry and the analysis based on the compound symmetry pattern displayed a much inflated false positive rate (supplementary figure 3). This suggests that the reported power gains mainly arise from increasing the de facto allowed false positive rate beyond the nominal significance level.

When comparing groups such as diseased vs healthy it is important to note that variances and correlations may differ between the groups. For instance, the protein abundances of healthy subjects may be more homogeneous and more predictable across repeated measurements corresponding to having smaller variances and stronger correlations than in the diseased population. Failure to account for differences in covariance parameters between the groups by including group-specific covariances in the linear mixed model, may lead to biased inferences, especially when the groups have different sizes [22].

### Hypothesis testing – t-tests vs F-tests and degrees of freedom methods

Statistical tests identify protein intensity profiles that vary across time or anatomical sites. Common approaches include:

i. Pairwise comparisons using ordinary paired t-tests
ii. F-tests based on linear mixed models
iii. Pairwise comparisons using t-tests based on linear mixed models

Pairwise comparisons offer an advantage over F-tests: while F-tests only detect whether a profile changes, pairwise tests can distinguish between different profiles, enabling interpretation of trajectory patterns thus providing valuable insight into the nature of the time evolution (or spatial differences). Importantly, making pairwise comparisons does not increase the false discovery rates despite the higher number of tests, since false discovery rates are *scalable* [23]; if the false discovery rate is controlled for each specific pair of time points (sites), it will also be controlled for the study overall. Comparisons should include all pairs of time point (sites) to avoid missing profiles with localized changes.

From a power perspective, pairwise comparisons generally outperform F-tests for profiles with localized changes, whereas F-tests may have higher power when changes develop gradually across all points. Between ordinary paired t-tests and linear mixed model based t-tests, results are equivalent under unstructured covariance and no missing data, but the linear mixed model handles missing data more optimally and enables inclusion of additional covariates [15].

A challenge with using linear mixed models is to compute the degrees of freedom for evaluating the t- and F-test statistics correctly in studies with a small to moderate number of participants. There are several methods for this with Satterthwaite and Kenward-Roger’s methods being the recommended choices [24, 25]. However, in our simulations neither approximation controlled the false positive rate at the nominal significance level when a conventional F-test was used for analysis (Supplementary Figure 2). We therefore suggest using a parametric bootstrap-version of the F-test to obtain valid statistical inference.

**Figure 2:**
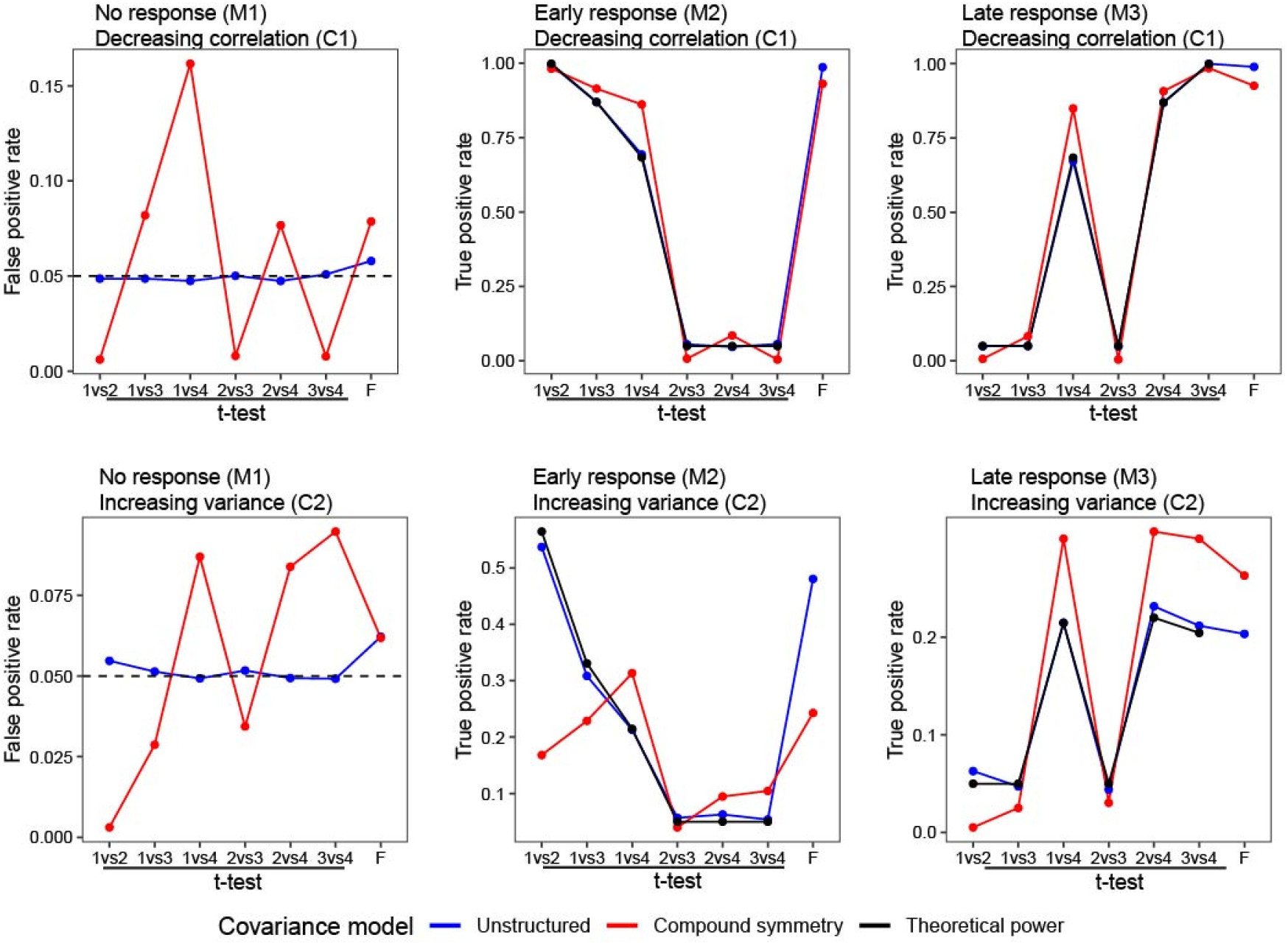
Benchmarking statistical models on simulated datasets. Four datasets were generated with different mean response and covariance structure as previously described. All datasets were analyzed with two statistical models assuming either unstructured covariance or compound symmetry. The left column shows the false positive rates from the analysis with the time-constant mean profile (M1) with either the C1 (top) or the C2 (bottom) covariance structure . The middle column shows the analysis of the early response profile (M2). The right column shows the similar analysis of the late response profile (M3). When basing the analysis on pairwise t-tests, the unstructured model (blue) achieves well-controlled false positive rates across all datasets, whereas compound symmetry (red) does not. The unstructured model also reproduces expected power (black), whereas the compound symmetry often fails due to misspecification of the covariance. Deviations between the F-tests based on compound symmetry and unstructured covariance are most pronounced for the data generated with covariance (C2) where the variance changes over time.

### Resources for Repeated Measures Analysis in R

There is no consensus software for analyzing repeated measures proteomics data, but several R packages provide useful functionality. Most approaches are based on linear mixed models with correlations modeled via subject-specific random effects.

The often-used LIMMA package [19], developed for the analysis of transcriptomics data, makes strong assumptions about the covariance, effectively assuming compound symmetry with constant correlations and variances across all measurement points and across all proteins. While this may be acceptable for small datasets, it is generally overly restrictive. The DREAM package [18] relies on an empirical Bayesian approach similar to LIMMA, but uses a more flexible approach, which estimates a new covariance structure for each transcript using random effects.

The packages timeOmics [17] and MSstats [26, 27] were specifically developed for proteomics data. Both approaches provide an analysis of proteome changes using linear mixed-effects models. The first uses splines to model the response over time, whereas the second is treating time as a categorical variable. Both model covariance via subject-specific random effects.

Covariance pattern models with flexible covariance structures are implemented in R-packages nlme [28], mmrm [29], and LMMstar [30]. The packages differ slightly in their selection of covariance patterns, but all include unstructured covariance. However, nlme does not make small sample adjustment to the degrees of freedom while mmrm has an option for using Kenward-Roger degrees of freedom and LMMstar by default uses the Satterthwaite approximation. Table 1 summarizes repeated measures models available in different R packages.

**Table 1.**
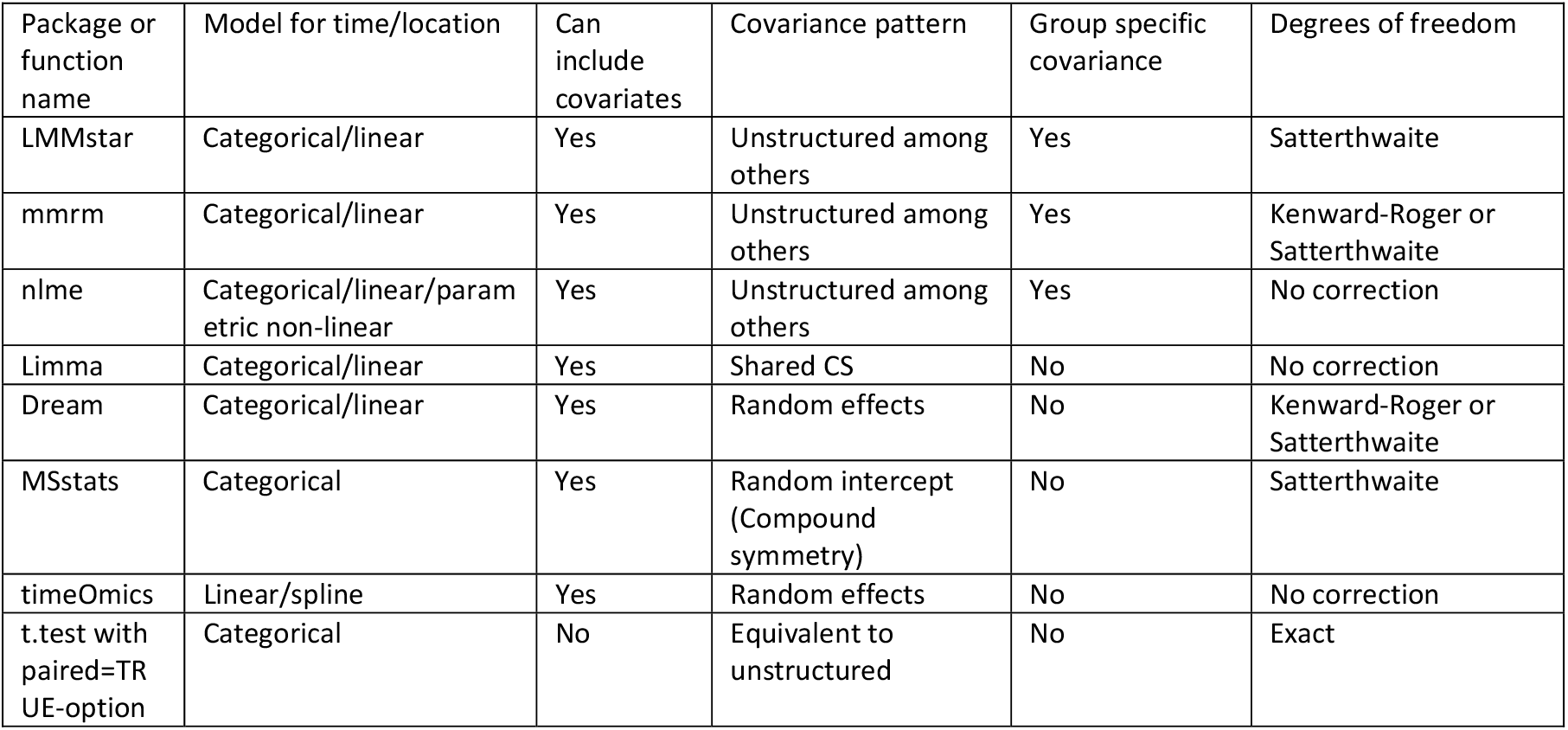
Summary of repeated measures models across R packages.

### Evaluation of statistical models

To evaluate the validity of a statistical test, we assess two key properties: control of the false positive rate and statistical power. The false positive rate is not necessarily controlled by the nominal (predefined) significance level due to deviations from model assumptions and the approximate nature of many tests. It is therefore important to evaluate a test and confirm that it maintains the nominal level (typically 5%). The power (true positive rate) of a statistical test reflects its ability to detect changes in the mean response over time or location. Once the false positive rate is well controlled, it is desirable to obtain the highest possible power.

To evaluate the performance of commonly applied statistical tests, we generated repeated measures datasets from varying multivariate normal distributions including mean-trajectories that either remained constant or changed over time. To simplify, two different time-changing patterns were considered: Early change (increase from baseline at timepoint 1 to a higher level maintained at timepoints 2-4) and Late change (constant mean from timepoints 1-3, followed by an increase at timepoint 4):

- M1 (time constant profile): µ1 = µ2 = µ3 = µ4 = 1
- M2 (Early change): µ1 = 1, µ2 = µ3 = µ4 = 1.5
- M3 (Late change): µ1 = µ2 = µ3 = 1, µ4 = 1.5

Covariance structures were varied by independently modeling the variance or correlation pattern as either decaying correlation over time or increasing variance across timepoints:

- C1 (standard deviations constant) σ1=σ2=σ3=σ4=1, (correlation decreasing over time) ρ12= ρ23= ρ34=0.8, ρ13=ρ24=0.5, ρ14=0.2
- C2 (standard deviations increasing over time) σ1=1, σ2=5/3, σ3=7/3, σ4=3, (constant correlation over time) ρ12 = ρ23 = ρ34 = ρ13 = ρ24 = ρ14 = 0.5

Illustrations of the resulting datasets are shown in Supplementary Figure 2. Initially, we simulated 10,000 datasets with n=40 subjects for each combination of mean profile and covariance. These were analyzed using the lmm-function in the LMMstar-package with the covariance pattern set to either compound symmetry or unstructured. We evaluated changes in mean using both pairwise t-tests and F-tests with degrees of freedom estimated by the Satterthwaite approximation (default in lmm).

First, we evaluated the effective significance level of the tests by computing the false positive rate in the datasets with mean profile constant over time. A valid statistical test should have an effective significance level which is less than or equal to the nominal 5% significance level. Secondly, we evaluated the power of the tests by computing the true positive rate in the datasets with profiles changing over time. Due to the poor small-sample performance of the conventional F-test, further simulations were made to compare the p-values obtained by means of the Satterthwaite approximation to those based on parametric bootstrapping. Details on the parametric bootstrap can be found in the Supplementary Material.

## 3. Results

Unstructured covariance model provides more reliable inference than compound symmetry

To assess the effect of both the choice of covariance pattern in the statistical analysis and the choice of statistical test, we analyzed the four different generated datasets with either an unstructured covariance structure or compound symmetry and using either t-tests or F-tests for comparisons. Figure 2 summarizes the false positive rate and power for both choices of covariance across all tests. The left column shows the analysis of the no response (M1 shape), which demonstrates the ability of the analysis to control the false positive rate. The unstructured covariance pattern is substantially more robust than the compound symmetry. The middle column shows the power of the two statistical models when analyzing the trajectories with early response (M2). The right column shows the analysis of the late response (M3) datasets. Additional comparisons to DREAM and Limma, which are based on mixed models with random effects, are presented in the Supplementary Figure 4.

The pairwise t-tests based on the unstructured covariance pattern correctly controls the false positive rate at the 5% significance level, whereas the compound symmetry pattern does not. This discrepancy arises because the compound symmetry model cannot capture the time-dependent changes in correlations and variances. Both versions of the F-test display false positive rates close to the nominal significance levels, albeit both are somewhat liberal with an effective significance level of 6-9% across all simulated datasets. The F-test associated with the compound symmetry model appears to be more robust to the misspecification of the covariance pattern compared to the pairwise t-tests, indicating that the bias in the estimated correlations and variances cancels across timepoints.

The true positive rates (bottom panel) show that the unstructured covariance pattern model matches the expected power for the paired t-tests, which can be deduced from the data generating model parameters. The t-tests based on the compound symmetry pattern have higher power for pairs of timepoints where the effective significance level is higher than the nominal 5% level and lower power for pairs of timepoints where the effective significance level is lower than the nominal 5% level.

In summary, the unstructured covariance is superior to compound symmetry both in controlling the false positive rate and in achieving the expected statistical power due to its ability to correctly capture time-varying correlations and variances. The simulations confirmed the poor small-sample performance of the F-test. Although the Satterthwaite approximation was used and 40 subjects is a fairly large sample size for a proteomics study, only the pairwise t-test maintained the nominal 5% significance level whereas the F-test was too liberal under both models. Given that the typical proteomics datasets is within a range of tens to a few hundred subjects, we address this further in the following section.

### Parametric bootstrapping improves the small sample performance for the F-test

We proceeded to investigate the problem of validity of the analytical F-test by a simulation study. As shown by the simulations in Figure 3, the F-test is too liberal when sample sizes are small. Because F-tests are widely used in repeated measures proteomics, this poses a general problem: p-values arebiased downwards for small cohorts, even when applying Satterthwaite or Kenward-Roger corrections (Supplementary Figure 2). We set out to evaluate the performance of various types of F-tests across a range of samples sizes common in proteomics studies. Specifically, we evaluated two datasets with early changes (M2) and either increasing variance (C2) or decreasing correlation (C1) with repeated measurements from 10 up to 40 subjects. Figure 3 shows the effective significance level of the F-tests based on the unstructured covariance pattern and the compound symmetry pattern with p-values calculated using either the Satterthwaite approximation or parametric bootstrapping.

**Figure 3:**
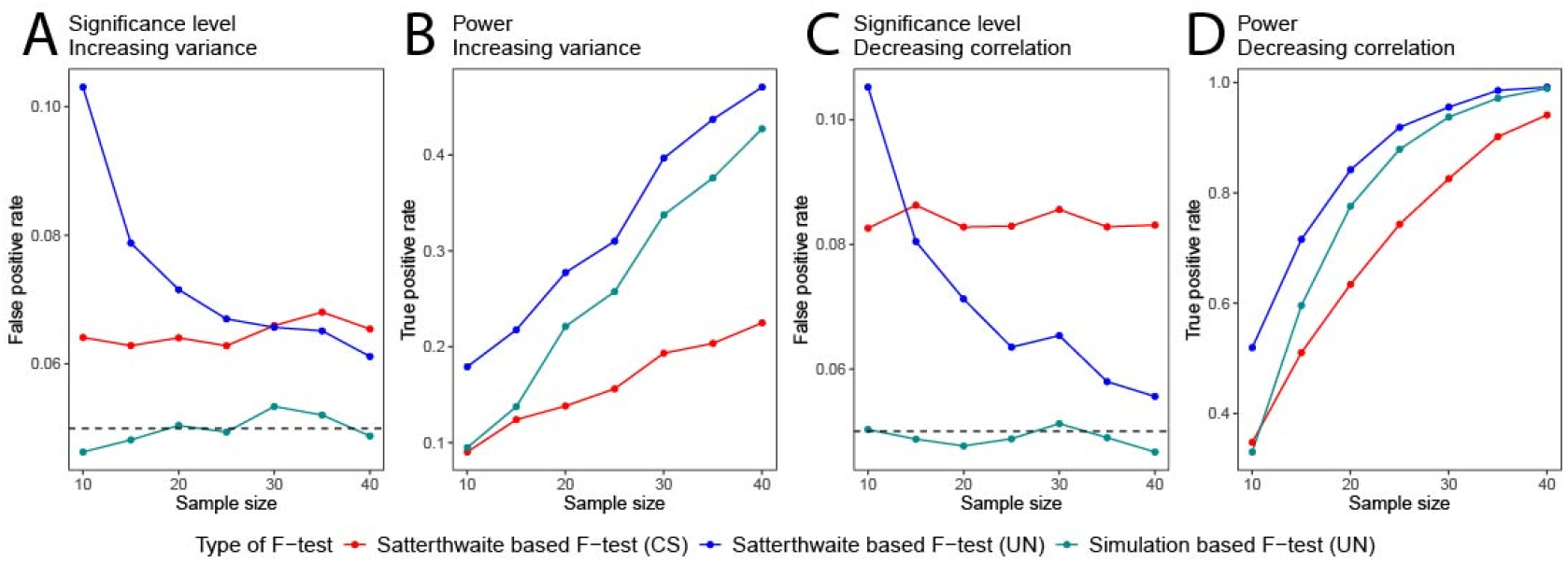
Effective significance level and power of conventional F-tests compared to parametric bootstrap tests as a function of sample size. Analyses are shown for simulated datasets with (A, B) early change (M2) and increasing variance (C2) and (C, D) early change (M2) and decreasing correlation (C1). Three different F-tests are compared, a compound symmetry covariance Satterthwaite F-test (red), an unstructured covariance Satterthwaite F-test (blue), and an unstructured covariance parametric bootstrap F-test (cyan). The conventional F-tests with degrees of freedom computed using the Satterthwaite approximation performs poorly even for moderately large sample sizes, while the parametric bootstrap test performs better, achieving a false positive rate of approximately 0.05. For smaller sample sizes, this reduces power, but appropriately so, as the test becomes more valid. The analysis based on the compound symmetry pattern is less biased for smaller sample sizes, compared to the unstructured covariance, but still has inflated false positive rates.

For the unstructured model, the false positive rate exceeds 0.10 for both datasets when only 10 subjects are included, far above the nominal 0.05 threshold, leading to a substantial inflation in false discoveries (Fig. 3A,C). The compound symmetry model is somewhat more robust at smaller sample sizes, for the C2 covariance structure (increasing variance) (Fig 3A), but not for the C1 covariance structure (decreasing correlation) (Fig 3C). Both analytical F-tests consistently have false positive rates above 0.05 (Fig 3A,C). To address this problem, we implemented a parametric bootstrap to evaluate the p-values. Specifically, we compute the p-value by comparing the obtained F-value to a null-distribution of F-values obtained by sampling from the best-fitting model satisfying the null hypothesis (see Supplementary Material). This substantially improves the control of the false positive rate (Fig. 3A,C), with only a minor reduction in power (Fig. 3B,D). While these simulations are based on idealized datasets and results will vary with real-world data characteristics, they underscore the importance of assessing the validity of the statistical test of choice. Our findings demonstrate that the parametric bootstrap F-test provides a practical solution for achieving accurate significance levels in small to moderate size repeated measures proteomics studies.

### T-test maintains accurate control of the false positive rate and improves power for detecting time-changing trajectories

The pairwise t-tests associated with the linear mixed model analysis present a compelling alternative to the F-test and will in many cases be a more robust alternative, despite the more widely use of F-tests in repeated measures proteomics. The issue with p-values that are biased downwards for small to moderate samples is specific to the F-test and do not affect the t-test, as shown in Figure 2. To assess this observation further, Figure 4 shows the performance of the t-tests based on the unstructured covariance on simulated data with the M2 response (early change), C2 covariance structure (increasing variance) and varying sample sizes. The false positive rate for all six possible comparisons between timepoints are shown. These results illustrate that the t-test maintains a well-controlled effective significance level (blue line) even with as few as 10 subjects, while the power (red line) increases as sample size grows. As expected, power decreases with increasing distance in time due to rising variance, but the effective significance level remains stable across sample sizes. This highlights that, unlike the F-test, the t-test is unaffected by small sample bias. Our findings indicate that the t-test is a reliable and often more powerful alternative to the F-test for identifying time-varying or site-dependent trajectories in repeated measures proteomics, particularly when sample sizes are limited.

**Figure 4:**
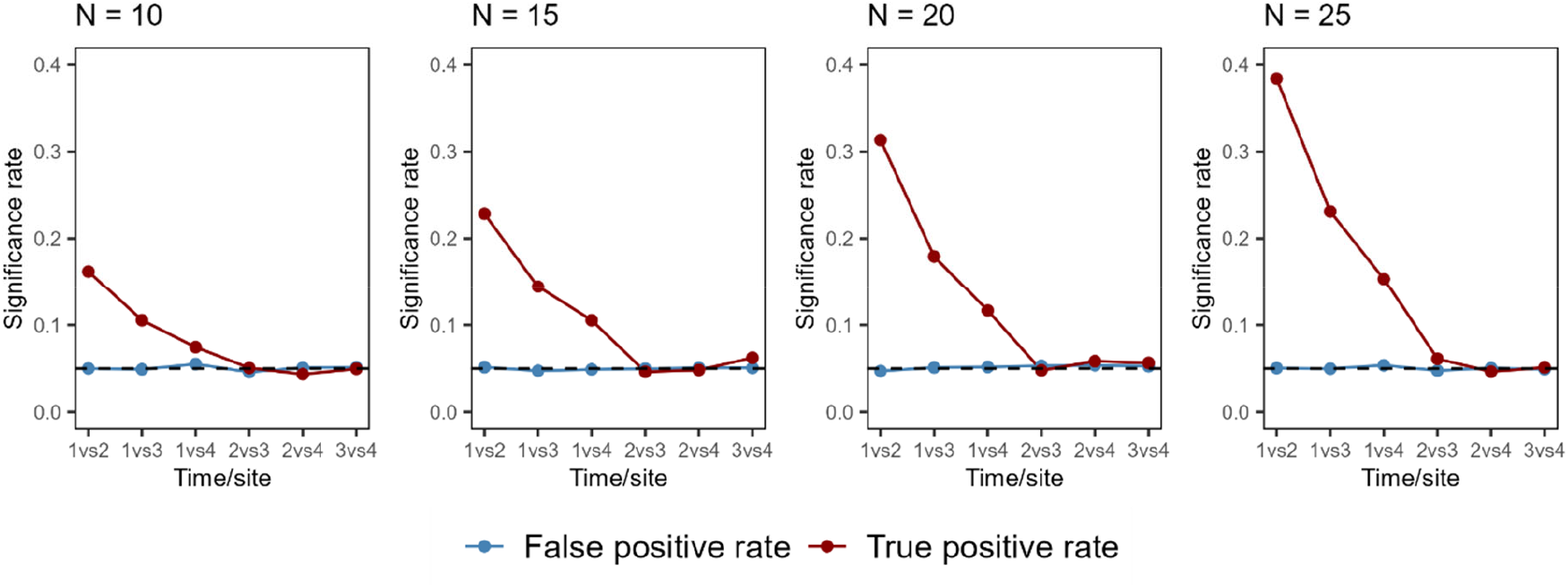
False positive rate and power of the pairwise t-test based on a linear mixed model with unstructured covariance pattern. Simulations were performed on a repeated measures dataset with four timepoints, M2 response and C2 covariance structure and sample sizes ranging from 10 to 25 subjects. The dashed black line marks the nominal significance level of 0.05. Results show that the t-test maintains a stable false positive rate (blue line) across all sample sizes, while power (red line) increases with larger cohorts. Only comparisons which include the first timepoint yield significant hits, due to the M2 response shape. The decline in power from 1vs2 to 1vs3 and 1vs4 reflects the increase in variance associated with the C2 structure.

## 4. Recommendations for the analysis of balanced repeated measures proteomic data

This section outlines a practical workflow for analyzing balanced repeated measures proteomics data, focusing on steps that differ from standard analyses of independent measurements. A schematic overview of these steps is shown in Figure 5. We assume that the proteomic data has been processed, normalized and quality-controlled prior to analysis, resulting in a matrix of values where rows represent proteins and columns represent samples. The values are estimates of protein abundance, such as protein intensities derived from the precursor intensities measured by the mass spectrometer. While this workflow is illustrated for mass spectrometry-based data, it generalizes to other proteomic platforms including affinity-based approaches [31, 32].

**Figure 5:**
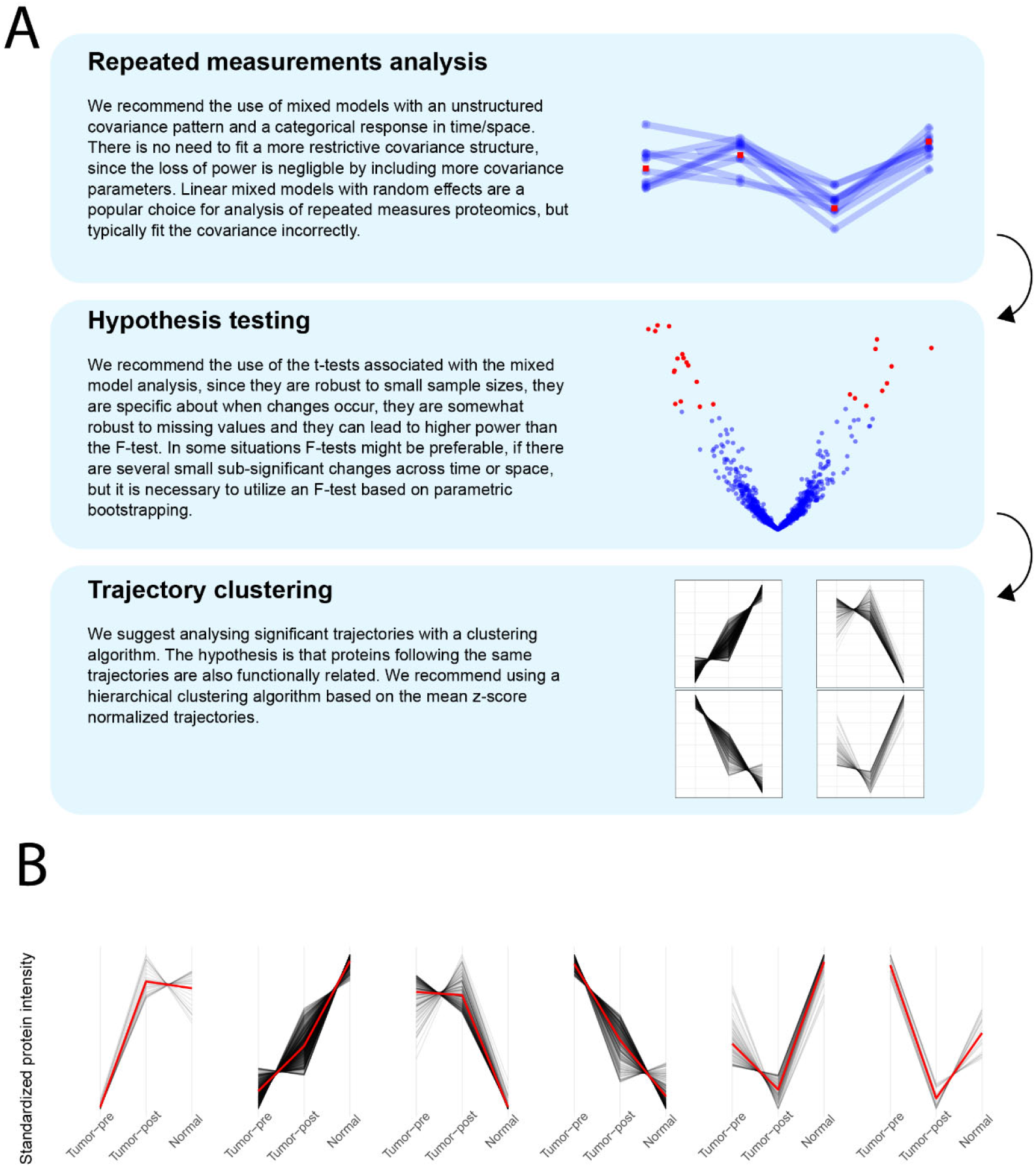
Workflow and results from a real-world repeated measures proteomics analysis. (A) Recommended workflow for analyzing repeated measures proteomics data. The key distinction from analyses of independent measurements lies in selecting an appropriate linear mixed model and corresponding hypothesis test to ensure control of the false discovery rate. Trajectory clustering is suggested as an optional downstream step for exploring patterns among significant trajectories. (B) Trajectory clusters of different response patterns of protein intensities in tumor tissue measured before treatment (Tumor-pre), tumor post treatment (Tumor-post), and in normal tissue post treatment in n=34 patients with breast cancer. Differences in protein intensities between time point and tissue type were identified using a linear mixed model with an unstructured covariance pattern. Full details of this analysis are provided in the supplementary material and the online repository.

### Repeated measurements analysis

The defining feature of a repeated measures analysis is the use of linear mixed models that account for correlated observations and potential variance heterogeneity. We recommend covariance pattern models with an unstructured covariance structure, which can be analyzed in R using either LMMstar or mmrm which include Satterthwaite or Kenward-Rogers correction to the degrees of freedom. We used the LMMstar package, but mmrm provides similar results (data not shown).

Specifying an appropriate covariance structure is critical. We recommend an unstructured covariance pattern, because it controls the risk of false positives regardless of time-dependent variance and correlation. For studies with multiple groups, estimating group specific unstructured covariance matrices can be beneficial, as variances and correlations may differ between diseased and healthy controls.

Outliers can strongly influence both estimated fixed effects and covariances. Since the goal is a robust estimate of the population level profile, single outliers should not determine which trajectories are significant. Outliers can e.g. be detected using studentized residuals to identify the measurements that deviate the most from the estimated population mean and scaled residuals to identify subjects who display highly unexpected changes relative to previous measurements (both are available from the LMMstar package). We recommend removing proteins with excessive missing values to ensure estimates that represent the broader population.

### Hypothesis testing and false discovery

The aim of the repeated measurements analysis is to identify proteins that change substantially over time (or space) and the time-intervals over which the change occurs. We recommend using pairwise t-tests based on linear mixed model analysis for hypothesis testing. All possible comparisons of time-points should be included with the false discovery rate controlled for each comparison separately. Using the t-tests for making pairwise comparisons, a trajectory is deemed substantially changing over a particular time-interval if the corresponding t-test produces a p-value smaller than 0.05 after FDR-adjustment. If the F-test is preferred to the t-test, we recommend that the p-value is evaluated using parametric bootstrapping as analytical approximations are insufficient even for moderately large sample sizes.

### Trajectory clustering

After identifying significant time- or location-dependent changes, clustering can provide an overview of the distinct patterns of change. We recommend hierarchical clustering based on mean standardized trajectories rather than sample-to-sample correlations, which may reflect unrelated effects. We used the pheatmap function in R for clustering and estimated the number of clusters using the fviz_nbclust plot from the factoextra library, though the number of clusters remains subjective.

### Downstream analysis

When clusters are identified, biological interpretation can begin, e.g. by distinguishing early versus late changes or spatial differences. Downstream analysis may include geneset enrichment or protein-protein interaction network, to decipher biological function of subclusters.

A detailed walkthrough of two real-world repeated measures proteomics datasets (“Weight Loss data” [7] and “Tumor data” [8]) is available in our online repository [33]. The trajectory clusters from both analyses are included in the Supplementary Material (Supplementary Figure 5 and 7), and trajectory clusters from the Tumor data are shown in Figure 5B. We additionally compared different linear mixed models using the Tumor data and evaluated their performance across a range of sample sizes by analyzing subsets of the original dataset. This comparison demonstrates that the choice of linear mixed model can substantially affect the number of proteins identified as significant (Supplementary Figure 6).

In summary, our recommended workflow utilizes a linear mixed-model with an unstructured covariance pattern to make pairwise comparisons with t-tests to identify proteins that change substantially over time (or space) followed by clustering of standardized mean trajectories. This approach ensures valid control of the false discovery rate and interpretable biological insights in repeated measures proteomics studies.

## 5. Discussion

The aim of this study was to provide robust strategies for analyzing repeated measures proteomics data. While statistical software for repeated measurements analysis exists and can be applied to proteomics (see Table 1) there is still no widely accepted standard approach. Repeated measurements analysis is inherently more complex than analysis of independent measurements and often relies on approximations. Its complexity warrants careful consideration of the analytical strategy to properly control the risk of false discoveries and optimize statistical power.

In the context of proteomics, we recommend the use of covariance pattern models, which are more flexible in capturing time- or site-varying variances and correlations likely to occur in biological data. It is beneficial to use a flexible model for proteomics analysis since it needs to adequately fit thousands of protein trajectories. Through simulation studies we have illustrated that the unstructured covariance pattern model is superior in fitting protein-specific correlation and variance structures leading to adequate control of the false positive rate and more optimal statistical power. In contrast, other more restrictive models such as linear mixed-effects models with correlation induced by random effects display inflated false positive rates or suboptimal statistical power whenever the restrictions fail to hold for specific proteins (other covariance patterns of complexity less than the unstructured covariance and greater than compound symmetry are discussed in [15]). Although model selection criteria such as AIC [15] or likelihood ratio tests [17] are often recommended, they can likewise lead to increased false positive rates since the “best” model out of a set is always chosen, which can lead to more spurious good fits. Our findings suggest that the unstructured covariance model is preferable, as it can reproduce any covariance structure found in real life data without a notable loss of power.

Based on our simulations, we further recommend using pairwise t-tests based on the linear mixed model with unstructured covariance. T-tests have a better small sample performance and provide interpretable information about when changes occur compared to the often-used F-tests. Our simulations revealed that with four repeated measurements for each of up to 40 subjects, the F-test has a false positive rate larger than the nominal 5% significance level, even when recommended small-sample adjustment to the degrees of freedom such as Satterthwaite and Kenward-Roger are employed. While a comprehensive evaluation of these violations is beyond the scope of this work, we recommend that researchers perform simulation studies of their intended statistical analyses, with the caveat that it is necessary to simulate data with heterogeneous covariance structures.

Power calculations are standard in experimental design, but we propose extending this practice to evaluations of the false positive rate to ensure nominal control.

We further demonstrated that the inflated false positive rate of the conventional F-test can be avoided by using parametric bootstrapping to evaluate the p-value. However, this comes at the cost of a substantial increase in computation time.

To illustrate our recommendations, we provided detailed analysis of two real-world datasets in the Supplementary material and a complete step-by-step workflow in a public repository [33]. This repository includes all simulations and practical guidance for implementing repeated measures analysis in time and/or space.

The surge of large-scale proteomics has led to increasingly complex study designs. This work provides practical recommendations for valid statistical analysis, ensuring control of false discoveries and proper representation of correlation and variance structures. Applying these strategies will enable more confident interpretation of biological mechanisms and disease trajectories, ultimately supporting robust and reproducible discoveries in proteomics.

## Acknowledgements

This work was supported by grants from Independent Research Fund Denmark (10.46540/5281-00023B) and the Novo Nordisk Foundation (NNF25OC0097951) to A.L. and fellowship funding from the Danish Cardiovascular Academy, which is funded by the Novo Nordisk Foundation (NNF20SA0067242) and The Danish Heart Foundation to M.S.S (PD2-2024003-DCA).

## Data Availability Statement

No new experimental mass spectrometry data were generated in this study. The real-world proteomics datasets analyzed in this work were obtained from publicly available supplementary materials accompanying the original publications by Geyer et al. [7] and Shenoy et al. [8]. Specifically, the datasets are available in Supplementary Table EV8 of Geyer et al. and Supplementary Table EV2 of Shenoy et al. The code used to generate the in silico datasets and perform all analyses is publicly available in the GitHub repository [33]. All simulation parameters required to reproduce the results are described in the manuscript and implemented in the deposited code.

