## Supplementary material for "A Statistical Guideline for Analyzing Repeated-Measures Proteomics Data"

### Fixed Effects: Modeling Population Level Effects of Covariates

Supplementary Figure 1 illustrates some of the different protein intensity profiles that can occur in a study with four follow-up times (sites). This is to illustrate the need for a flexible statistical model.

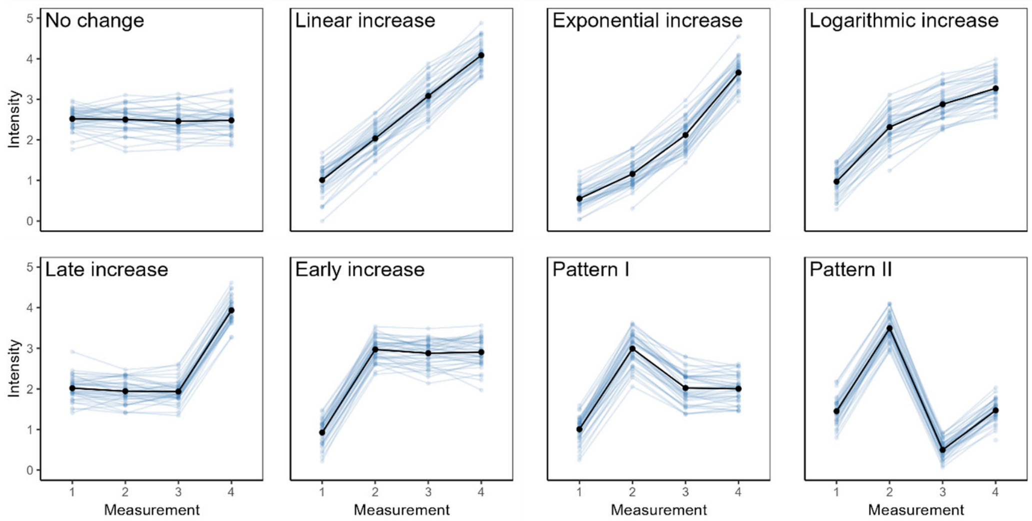


***Supplementary Figure 1:*** *Modeling time or site as a categorial fixed effect in a linear mixed model is essential to capture diverse protein intensity profiles*. Simulated data illustrating eight distinct protein intensity profiles across four repeated measurements (1-4). Estimated population means from a linear mixed model (black) are superimposed on observed data (blue). Modeling time (or anatomical site) as categorial avoids restrictive assumptions such as linearity, which can bias results and fail to reflect complex biological patterns. In these examples, with complete data and no additional covariates, fixed-effect estimates coincide with the sample means.

### Illustration of simulated datasets

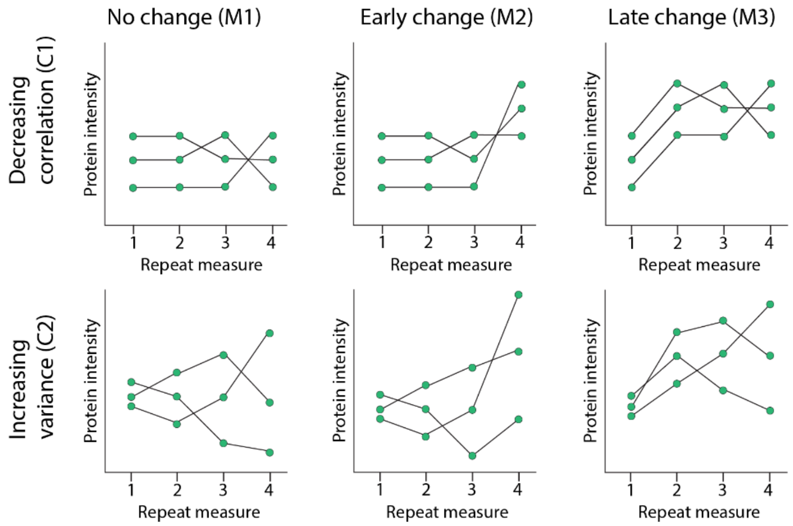


***Supplementary Figure 2:*** *illustration of the simulated datasets used in the main article*. All datasets are based on either of two covariance structures: decreasing correlation (C1) or increasing variance (C2), with either one of three response shapes: No change (M1), Early change (M2), or Late change (M3).

### Analytical small sample corrections are not sufficient for small to midrange sample sizes: Kenward-Roger versus Satterthwaite

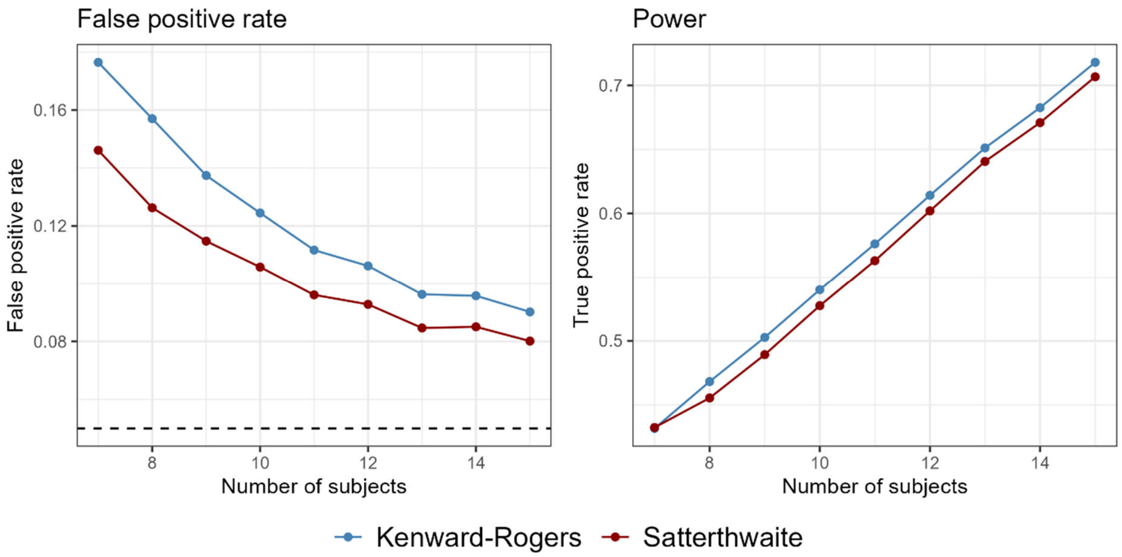


***Supplementary figure 2***. *Comparing analytical corrections for small sample sizes: Kenward-Roger versus Satterthwaite. We have compared the two approaches for a single in-silico simulated dataset with variable number of subjects. The regressions were performed using the mmrm package and trajectories were deemed significant according to an analytical F-test. The Kenward-Roger approach is slightly better for very small sample sizes, but both corrections are very far from the nominal significance level at 5%.*

The hypothesis tests associated with mixed model regressions are known to be imprecise for smaller sample sizes. Two canonical corrections have been developed to arrive at better estimates for the degrees of freedom: the Kenward-Roger[1] and Satterthwaite[2]. We analyzed simulated data with responses M1 (no change) and M2 (early change) and with the C1 covariance structure (decreasing correlation), as specified in the main manuscript. As shown in Supplementary Figure 2, both the Kenward-Roger and the Satterthwaite correction result in an inflated level of false positives compared to the nominal level of 0.05. For these simulations, Satterthwaite outperforms Kenward-Rogers in controlling the false positive rate. However, a previous in silico study found that Kenward-Roger degrees of freedom performed better than Satterthwaite in a randomized baseline follow-up design with missing data [3]. It is beyond the scope of this article to map out the differences between the Kenward-Roger and Satterthwaite corrections, especially since both these analytical approaches are too imprecise for a proper proteomics analysis.

### Relation between the number of covariance parameters and statistical power

In the context of covariance pattern models, there is a substantial difference between the number of parameters estimated for unstructured models (proportional to $n^2$) versus compound symmetry models (only two parameters). If the number of covariance parameters affects the power, the unstructured model will arrive at a lower power, if it does not capture more complexity than the compound symmetry model. To test this, we used both model types to analyse in silico-generated data with a compound symmetry structure, shown in Supplementary Figure 2. This should demonstrate the isolated loss of power from estimating more parameters, since the data complexity can be captured entirely by a compound symmetry model. We also included a dataset with a deviance from the compound symmetry structure at the last time point. The covariance structures are either “low” or “high” correlation, and either a fully compound symmetry structure or with a small perturbation as follows:

$$\Sigma_{const}^{\text{low}} = \begin{pmatrix} 1 & 0.5 & 0.5 & 0.5 \\ 0.5 & 1 & 0.5 & 0.5 \\ 0.5 & 0.5 & 1 & 0.5 \\ 0.5 & 0.5 & 0.5 & 1 \end{pmatrix} \Sigma_{var}^{\text{low}} = \begin{pmatrix} 1 & 0.5 & 0.5 & 0.2 \\ 0.5 & 1 & 0.5 & 0.5 \\ 0.5 & 0.5 & 1 & 0.5 \\ 0.2 & 0.5 & 0.5 & 1 \end{pmatrix}$$

$$\Sigma_{const}^{\text{high}} = \begin{pmatrix} 1 & 0.8 & 0.8 & 0.8 \\ 0.8 & 1 & 0.8 & 0.8 \\ 0.8 & 0.8 & 1 & 0.8 \\ 0.8 & 0.8 & 0.8 & 1 \end{pmatrix} \Sigma_{var}^{\text{high}} = \begin{pmatrix} 1 & 0.8 & 0.8 & 0.5 \\ 0.8 & 1 & 0.8 & 0.8 \\ 0.8 & 0.8 & 1 & 0.8 \\ 0.5 & 0.8 & 0.8 & 1 \end{pmatrix}$$

The response for the significant trajectories has mean values:

$$\mu_{low} = \begin{pmatrix} 1 & 1.4 & 1.4 & 1.4 \end{pmatrix}$$

$$\mu_{high} = \begin{pmatrix} 1 & 1.2 & 1.2 & 1.2 \end{pmatrix}$$

$$\mu_{ns} = \begin{pmatrix} 1 & 1 & 1 & 1 \end{pmatrix}$$

Supplementary Figure 2 shows a minor loss of statistical power associated with the unstructured model. The F-tests are performed using parametric bootstrapping. There is the most loss in power with the F-test associated with the unstructured model, when the data follows a perfect compound symmetry structure. Very little power is lost when using t-tests. However, with small deviations from the compound symmetry structure, the unstructured model is substantially more precise. There is little to gain from imposing a rigid covariance structure, but a lot to lose.

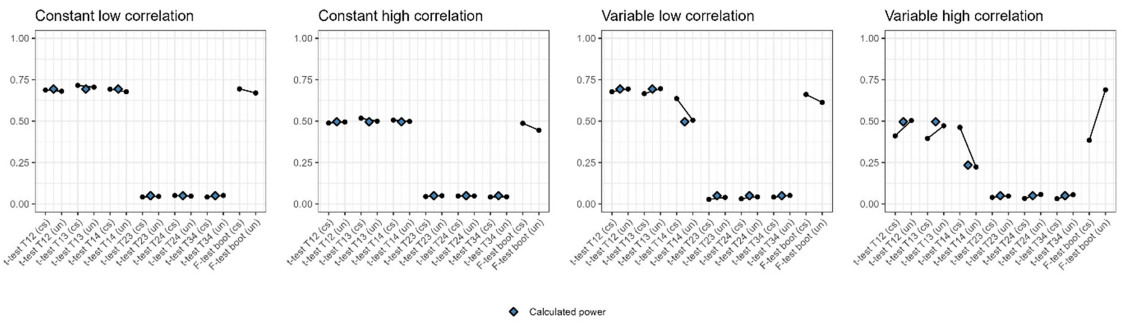


***Supplementary figure 3*** *Comparing power of statistical tests applied to data with various covariance structures. “cs” refers to compound symmetry and “un” refers to an unstructured covariance model. These simulation studies show that very little power is lost by using many covariance parameters, but a lot is lost by not being able to fit the data appropriately. The blue squares show the theoretical power for the t-test.*

### Performance by DREAM and LIMMA

We tested the F-test performance of DREAM and LIMMA on in silico generated datasets. The supplementary figure 5 shows the results for one data structure with covariance structure C1 (decreasing correlation) and early response (M2). Neither DREAM nor LIMMA has good significance level control and they are very similar to the covariance pattern model with the compound symmetry covariance structure. The F-test is analytical and corrected with Satterthwaite.

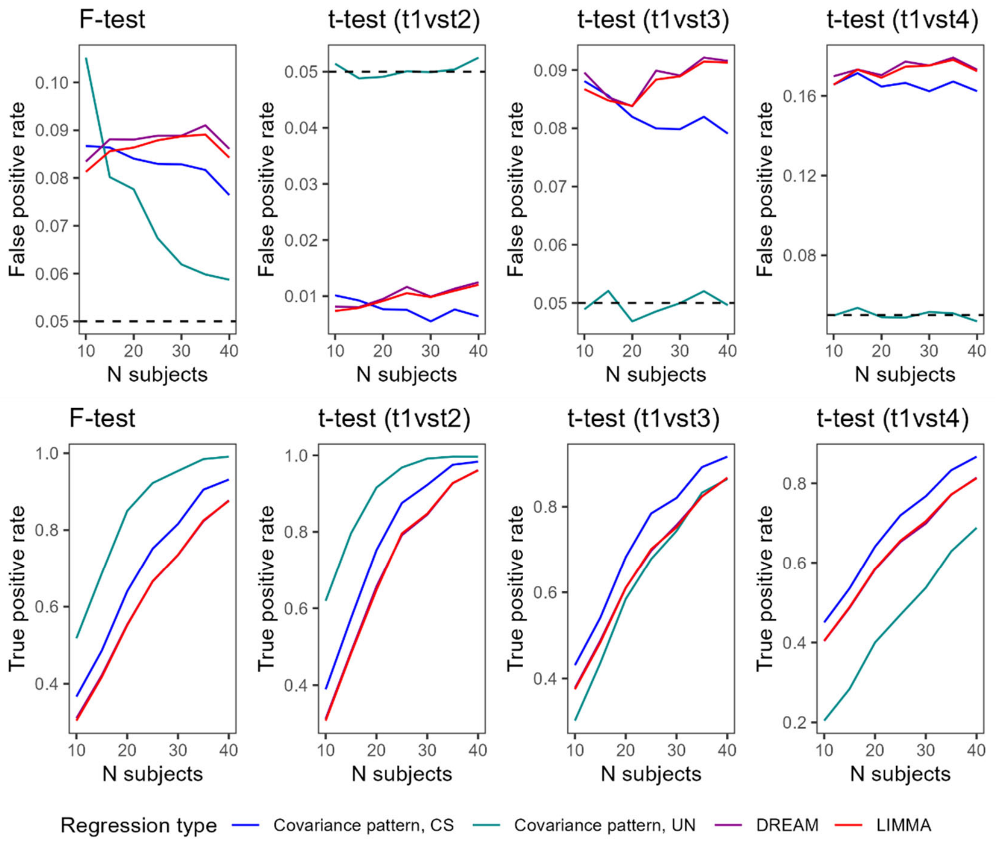


***Supplementary figure 4*** comparison covariance pattern models with either unstructured covariance or compound symmetry to analysis performed using LIMMA and DREAM with random intercepts. LIMMA and DREAM both behave like a compound symmetry, as expected with from the random intercept structure, and have difficulties in arriving at correctly controlled false positive rates. Results are for the F-test and three t-tests comparing timepoint 1 to timepoint 2,3 and 4. Only the unstructured covariance model arrives at correct false positive rates.

### F-test based on parametric bootstrapping

Since the analytical F-test is only approximate for repeated measures statistical models and handles small sample sizes poorly, we propose an F-test based on parametric bootstrapping. The method is as follows:

1) Fit a full regression model to the data and store the original F-statistic.
2) Fit a null-model to the data, with an intercept only in the response. Use the estimated response and covariance matrix from this fit to generate new datasets.
3) Repeat the following N times

a) Generate a new dataset the same size as the original, using a multivariate normal distribution with the response and covariance estimated in 2)
b) Fit this data with a full model with time as a covariate.
c) Store the F-statistic from this.

4) Finally, compare the original F-statistic to the generated distribution of F-statistics. The p-value is obtained by calculating how many F-values in the null hypothesis are higher than the original F-value.

### The power of a t-test

The power of t-tests associated with a mixed model fit are estimated using the base R function power.t.test(). However, there is a need to supply the pooled variance, to get a prober estimate of the power. The t-statistic is defined by the difference in mean divided by the pooled standard deviation. The pooled standard deviation is estimated from the formula for variation of a difference between two stochastic variables:

$$\sigma_{Y-X}^2 = \sigma_X^2 + \sigma_Y^2 - 2\sigma_X\sigma_Y r_{XY}$$

Where $r_{xy}$ is the Pearson correlation between the two variables.

### Tumor proteome study

Shenoy et al. investigated the change in tumor proteomes before and after chemotherapy, while also including post-treatment healthy tissue adjacent to the tumor [4]. This design contains repeated measurements across time and tissue type from the same patient. The study used paired t-tests to identify proteins changing significantly across conditions (normal tissue vs pre-therapy, normal tissue vs post-therapy, and pre-therapy vs post-therapy). In the main manuscript we discuss problems with the use of paired t-tests, instead of t-tests associated with a mixed model.

We re-analyzed the Tumor dataset by applying a mixed model with unstructured covariance pattern to each protein trajectory. The conditions were implemented in the statistical model as a categorical variable. Only trajectories with at least 70 percent non-missing values were included. To illustrate the ability of mixed models to include more covariates, we added the clinical variable “Lymph.node.involved” to the statistical model.

Supplementary Figure 6A shows a trajectory cluster analysis of all protein intensity trajectories deemed significant, where significance is defined by at least one out of the three T-tests having a false discovery rate value of 0.05 or less. The results of the clustering analysis illustrate the advantage of analyzing full trajectories in contrast to only focusing on pairwise comparisons. E.g. cluster 1 and cluster 2 both represent proteins with a substantial increase between pre-treatment tumor tissue and normal tissue. However, cluster 1 contains proteins that arrive at the same concentration as in normal tissue, whereas proteins in cluster 2 have intensities closer to the pre-treatment tumor than to the normal tissue. This could be quite informative in terms of determining which biological mechanisms are still delayed in normalizing following tumor treatment.

It is beyond the scope of this article to analyze the biological mechanisms of each trajectory cluster, but we have included an overrepresentation analysis of terms from the database GO

Biological Processes for illustrative purposes. Supplementary Figure 6B shows the three clusters with significantly overrepresented terms. We refer to the online material for the full analysis in R [5].

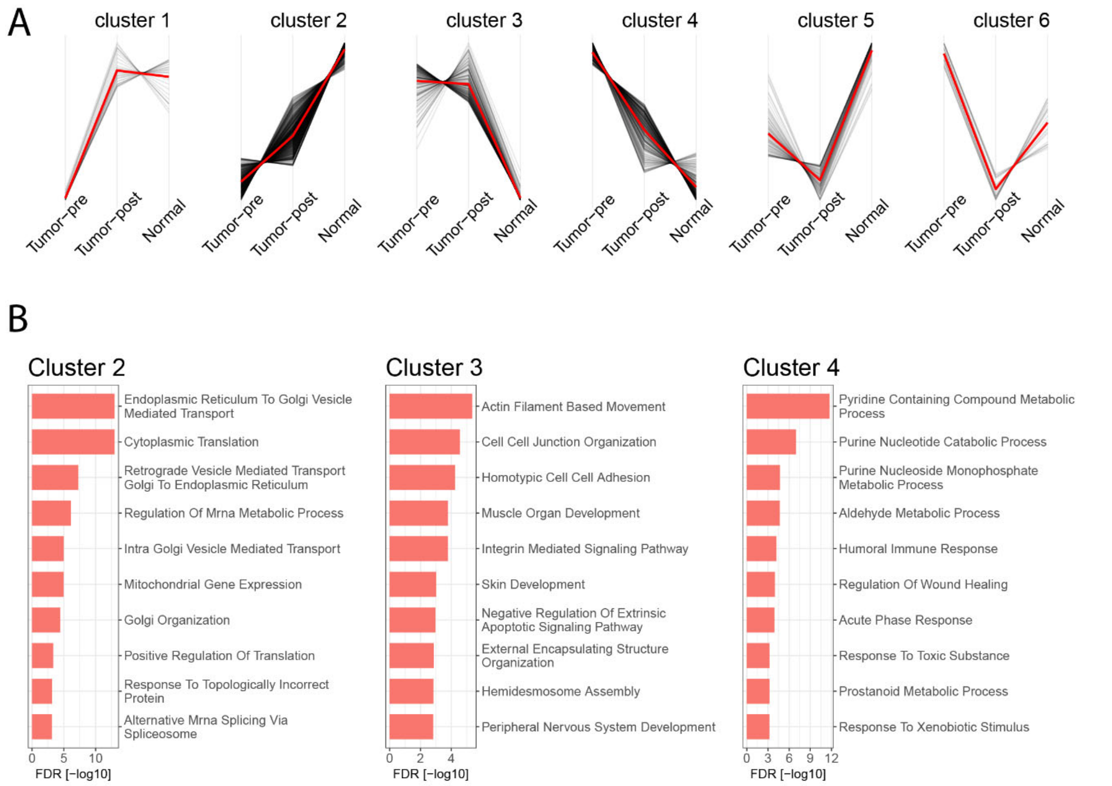


*Supplementary Figure 5. Regression analysis of Tumor proteome data to identify proteins changing significantly across conditions. (A) Clustering analysis of protein intensity trajectories across the three conditions. Each protein is represented as a line showing the mean of intensities standardized across all samples. Six clusters were chosen to represent the different trajectory shapes of the dataset. (B) Overrepresentation analysis of each trajectory cluster. The genesets are from the Gene Ontology Biological Processes database. Overrepresentation was tested using a hypergeometric test. The redundancy of genesets was additionally reduced using an in-house generated algorithm based on gene overlap between genesets. The algorithm is available online [5]. Three out of six clusters contained significant genesets.*

### The choice of statistical model has considerable impact on the results

Different versions of linear mixed models, with different covariance assumptions, arrive at a different number of significant hits. To illustrate the effect of model choice, we resampled from the Tumor dataset to construct datasets of varying sizes and analyzed the datasets with Limma, Dream, and an unstructured covariance pattern model. The resampling was repeated 5 times, and the mean results are presented in Figure 1, showing significant hits after Benjamini-Hochberg correction as a function of cohort size. Despite analyzing identical datasets, the statistical models led to different results. Since this is real-world data, there is no gold standard, but the choice of analysis led to different sets of significant proteins. In this dataset, three samples were obtained from each patient: pre-treatment tumor tissue, post-treatment tumor and tumor-adjacent normal tissue. The difference between pre-treatment and post-treatment tissue gave 247 hits with the unstructured covariance model, 142 hits with Dream and 63 hits with Limma, for the full patient cohort (34 patients). The pattern is reversed for the two other comparisons: pre-treatment vs tumor-adjacent and post-treatment vs tumor-adjacent. Here the unstructured covariance model was more conservative, whereas Dream produced an elevated number of significant hits, which likely reflects an inflation in false positives. The difference between models is especially pronounced for smaller cohort sizes in the post-treatment vs tumor-adjacent comparison: for 10 patients, the unstructured model arrived at 17 significant hits, Limma gave 65, whereas Dream arrived at 113. The unstructured covariance model is more flexible in representing the covariance structure of the data, so the difference in significant hits is likely related to a closer fit to the true covariance structure. The choice of statistical model can lead to an error-prone selection of significant protein hits, with both an increase in false negatives, leaving out important information, and false positives, clouding the biological interpretation with irrelevant proteins.

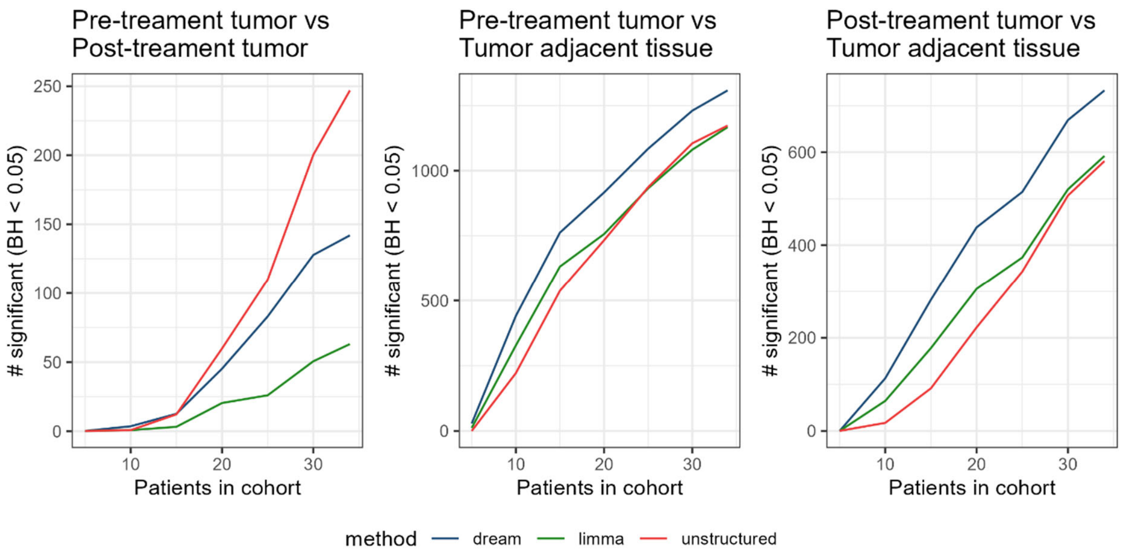


*Supplementary Figure 6. The number of significant proteins in the tumor dataset as a function of cohort size differs by statistical model. Analysis of the previously published Tumor dataset, with resampling at the patient level to construct datasets of varying cohort sizes ranging from 5 patients to the full cohort of 34 patients. Resampling was repeated 5 times, with lines representing the mean number of significant proteins (Benjamini-Hochberg corrected) as a function of cohort size. In this dataset, three samples were obtained from each patient: pre-treatment tumor tissue, post-treatment tumor tissue, and tumor-adjacent normal tissue. Data were re-analyzed using Limma, Dream, and a linear mixed model with unstructured covariance. The analysis is shown separately for each pairwise comparison of tissue type: pre-treatment vs. post-treatment, pre-treatment vs. tumor-adjacent, and post-treatment vs. tumor-adjacent.*

### Obesity proteome study

Geyer et al. investigated the proteomic changes in the plasma due to an 8-week weight loss intervention[6]. They have one control sample prior to the weight loss intervention (-8 weeks) and six samples after the weight loss (0, 4, 13, 26, 39, and 52 weeks), to follow changes in the proteome during weight maintenance. The study utilizes paired t-tests to detect changes between the baseline and any of the other time points. Multiple comparison correction is not applied directly, instead the study sets a nominal significance level at $5 \cdot 10^{-4}$, which identifies

84 proteins as significant. The study performs a hierarchical clustering algorithm based on standardized trajectories equivalent to our suggestion.

We re-analyzed the data using a mixed model with unstructured covariance pattern. All timepoints were compared to the -8 weeks control and trajectories were deemed significant if at least one t-test produced a false discovery rate equal to or below 0.05, using the Benjamini-Hochberg estimation. Our approach identified 178 significant trajectories, compared to the 84 proteins presented in the study. The significant trajectories were clustered and are presented in Supplementary Figure 7A. Supplementary Figure 7B shows the results of an overrepresentation analysis of gene sets from the GO Biological Processes database. Cluster 2 has an interesting adaptive shape, with proteins increasing during weight loss and then approximately decaying to the initial value; these include activation of an immune response.

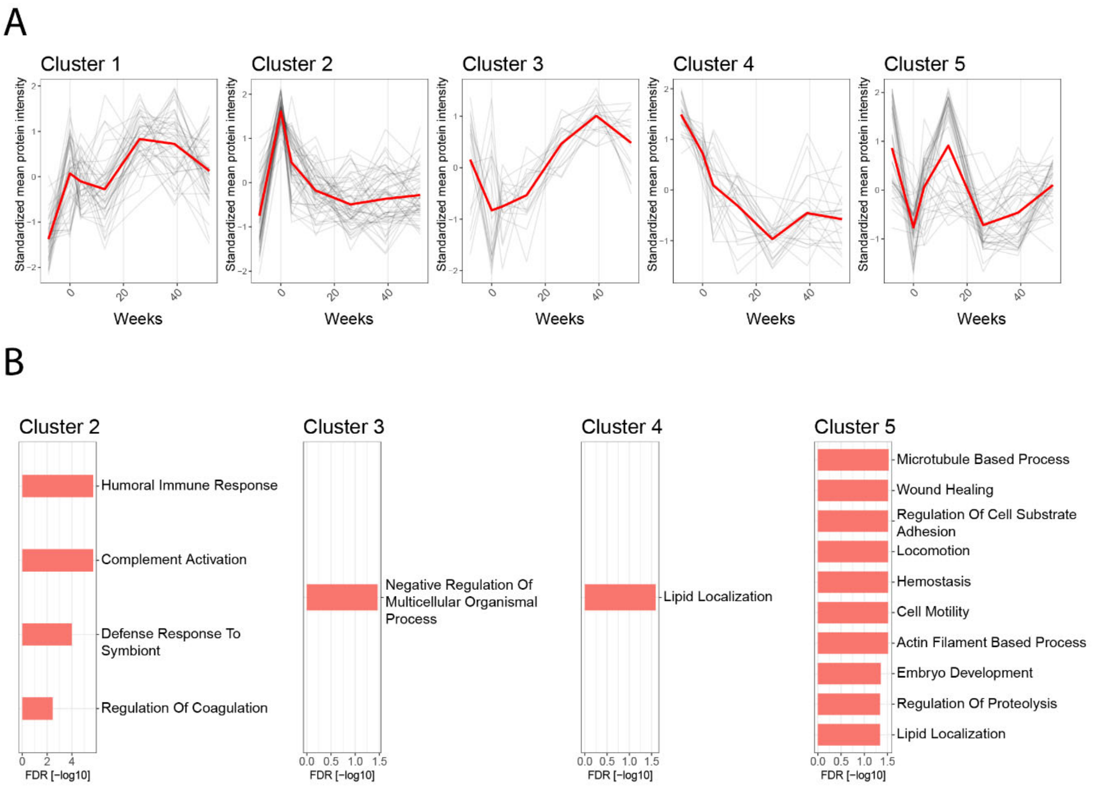


*Supplementary Figure 7. Regression analysis of Obesity proteome data. (A) Clustering analysis of protein intensity trajectories across the seven timepoints. Each protein is represented as a line showing the mean of intensities standardized across all samples. Five clusters were chosen to represent the different trajectory shapes of the dataset. (B) Overrepresentation analysis of each trajectory cluster. The gene sets are from the Gene Ontology Biological Processes database. Overrepresentation was tested using a hypergeometric test. The redundancy of gene sets was additionally reduced using an in-house generated algorithm based on gene overlap between gene sets. Four out of six clusters contained significant gene sets.*
